# Efficient propagation of misfolded tau between individual neurons occurs in absence of degeneration

**DOI:** 10.1101/372029

**Authors:** Grace I Hallinan, Mariana Vargas-Caballero, Jonathan West, Katrin Deinhardt

**Affiliations:** Biological Sciences, University of Southampton, UK; Faculty of Medicine and Centre for Hybrid Biodevices, Institute for Life Sciences, University of Southampton, UK

**Keywords:** Alzheimer’s disease, tau, aggregation, seeding, prion-like propagation, microfluidic devices

## Abstract

In Alzheimer’s disease, misfolded tau protein propagates through the brain in a prion-like manner along connected circuits. Tauopathy correlates with significant neuronal death, but the links between tau aggregation, propagation, neuronal dysfunction and death remain poorly understood, and the direct functional consequences for the neuron containing the tau aggregates are unclear. Here, by monitoring individual neurons within a minimal circuit, we demonstrate that misfolded tau efficiently spreads from presynaptic to postsynaptic neurons. Within postsynaptic cells, tau aggregates initially in distal axons, while proximal axons remain free of tau pathology. In the presence of tau aggregates neurons display axonal transport deficits, but remain viable and electrically competent. This shows that misfolded tau species are not immediately toxic to neurons, and suggests that propagation of misfolded tau is an early event in disease, occurring prior to neuronal dysfunction and cell death.

---

Neurofibrillary tangles (NFTs), a pathological hallmark of Alzheimer’s disease (AD), consist of insoluble aggregates of hyperphosphorylated tau protein. In AD, NFTs accumulate first in the entorhinal cortex, and with disease progression they appear in neighbouring regions such as the hippocampus, followed by the neocortex in later stages^1^. The neuroanatomical localisation of NFTs in AD brains suggests that tau pathology propagates through the brain along anterograde connected brain circuits^2–4^. Indeed, it is well established that pathogenic tau spreads between cells *in vitro*^5,6^ and *in vivo*^4,7,8^ in a prion-like manner, inducing the misfolding of native tau^5, 9–11^. Tau seeds, the tau species that template a misfolded conformation to native tau, are thought to be monomers or lower molecular weight oligomers rather than the high molecular weight tangles observed at late stages of disease, but their exact nature is still under debate^12–16^. Moreover, experiments using AD brain extracts to seed the misfolding of tau reported the existence of distinct pathological tau strains^6^. Pathogenic tau causes cell death both *in vivo*^4^ and *in vitro*^17,18^. Therefore, it has been suggested that one mechanism for the propagation of tau pathology is release following the disintegration and death of the tangle-bearing neurons^19^. In contrast, recent studies have shown that intact neurons exist in areas of pathology propagation in mouse models^20^, and have detected tau species capable of seeding misfolding in brain areas free of tangle pathology in human brains^21^. This suggests potential release of pathogenic tau from living neurons. However, loss of neurons is progressive within areas affected by degeneration, and as a consequence the *in vivo* studies were short of the resolution required to identify individual tau-propagating neurons.

In experimental paradigms, exogenous pre-formed recombinant tau fibrils or extracts from brains with tauopathy are added to neurons to investigate the mechanisms underlying tau pathology transmission. Studies using such exogenous tau preparations have shown that tau aggregates are internalised into primary neurons, can be trafficked both anterogradely and retrogradely along axons, and spread to and between connected cells^22–24^. However, these studies did not examine the functional outcome of tau misfolding, aggregation and propagation within neurons. In this study, we created a minimalistic neuronal circuit within a compartmentalised microfluidic device to investigate tau misfolding and propagation at single cell resolution. We show that a phosphomimetic tau, tau^E14^, in which 14 disease-relevant serine/threonine residues have been mutated to glutamate to mimic phosphorylation^25^, misfolds and aggregates in the absence of exogenous seeds. Misfolded tau^E14^ seeds template a rapid and efficient prion-like misfolding of native tau, and transmit a conformational change of tau between intact, connected neurons with high efficiency. This suggests that propagation of misfolded tau occurs between live, functioning neurons in very early stages of tau pathology prior to neuronal degeneration. In the context of AD, our findings imply that transmission of misfolded tau through the brain occurs between viable neurons and may precede detectable symptoms; it thus may present a disease modifying target for mild cognitive impairment.

## Results

### Phosphomimetic tau spontaneously aggregates in cultured neurons

Hyperphosphorylation of tau is associated with pathology. Pseudohyperphosphorylated tau (tau^E14^) mislocalises to dendritic spines and causes synaptic dysfunction^25^. To examine whether tau^E14^ spontaneously misfolds and oligomerises into aggregates such as observed in AD, we cultured murine hippocampal neurons and transfected them at day in vitro (DIV) 1 with fluorescently tagged pseudohyperphosphorylated or wild-type human 0N4R tau: GFP-tau^E14^ or RFP-tau^E14^ and GFP-tau^WT^ or RFP-tau^WT^, respectively. The use of GFP-tagged tau to study tau pathology propagation has recently been validated *in vivo*^26^. Expression of either tau^WT^ or tau^E14^ did not adversely affect axonal outgrowth when compared to control neurons (Fig. 1a). At DIV14, both GFP-tau^WT^ and RFP-tau^WT^ display a smooth and even distribution throughout the cell (Fig. 1b,d,e). In contrast, GFP-tau^E14^ and RFP-tau^E14^ expression resulted in a clustered tau distribution along distal axons (Fig. 1c,d,e, RFP not shown). Using an antibody that selectively recognises misfolded tau, MC1^27^, we confirmed that the clustering of fluorescence is reporting tau misfolding, which is not detected in axons exogenously expressing tau^WT^ (Fig. 1e). In order to analyse the appearance of tau aggregation within the axon we generated fluorescence distribution profiles along individual axons. This confirmed highly variable fluorescence intensities along axons of tau^E14^ expressing neurons, in comparison to a smooth fluorescence distribution in tau^WT^ expressing neurons (Fig.1d, S1c). Neurons were scored as positive for aggregation along the distal axon based on this differential variation (Fig. 1d, S1, supplemental methods). This analysis showed that at DIV14, 56.0 ± 7.7% of the tau^E14^ expressing axons have developed aggregates, whereas the distal axons of tau^WT^ expressing neurons remained aggregate-free (Fig. 1f). This demonstrates that pseudohyperphosphorylation of tau is sufficient to induce misfolding and aggregation of tau.

**Figure 1.**
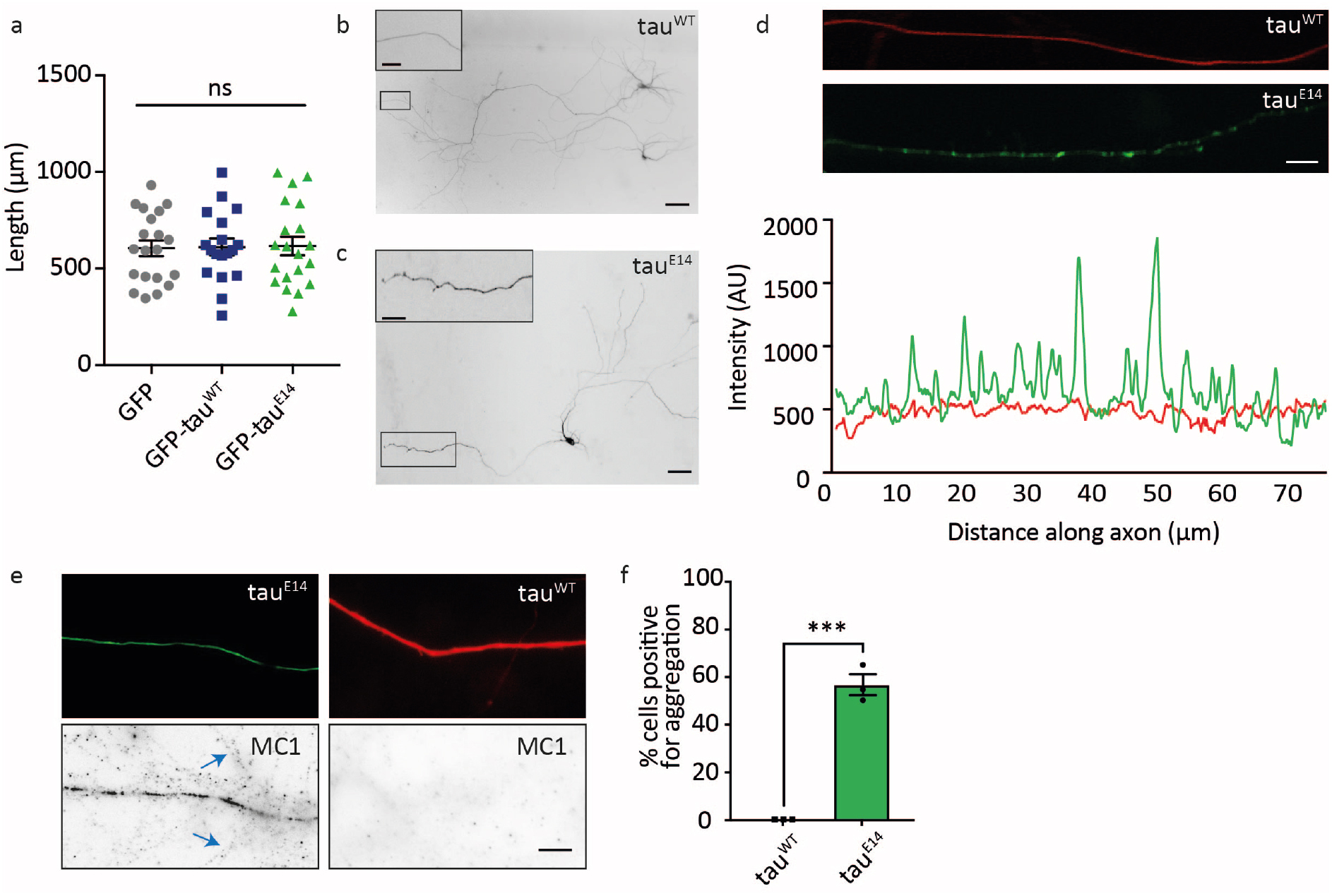
Tau aggregates spontaneously develop in GFP-tau^E14^ expressing hippocampal neurons.

(a) Hippocampal neurons were transfected with GFP, GFP-tau^WT^ or GFP-tau^E14^ at DIV1 and fixed and imaged at DIV7. No difference in axonal outgrowth was observed. Each data point is one axon, n≥18 axons per condition from 3 experiments. One-way ANOVA, p=0.9836, F_2,55_ = 0.01657. Error bars=SEM. (b, c) At DIV14, exogenously expressed tau localises throughout the neuron with (b) tau^WT^ outlining the microtubules and (c) tau^E14^ displaying aggregates in the distal axon. Scale bars 20 μm; insets, 5 μm. (d) A line was drawn along distal axons expressing tau^WT^ (top, red) or tau^E14^ (middle, green) to generate corresponding intensity profiles (bottom). Scale bar, 5 μm. (e) Tau^E14^, and surrounding axons (blue arrows), but not tau^WT^ expressing axons are positive for misfolded tau. Scale bar, 5 μm. (f) Analysis of fluorescence intensities along axons shows that at DIV14, aggregates selectively appear in a population of tau^E14^ expressing axons. Each data point is the mean of one experiment, n≥12 axons per experiment. Mann-Whitney U test, p<0.0001. Error bars=SEM.

### Tau aggregation efficiently propagates to connected neurons

MC1 staining to visualise misfolded tau revealed that neurites in the vicinity of a tau^E14^ expressing axon were positive for misfolded tau, whereas those surrounding an axon expressing tau^WT^ remained MC1 negative (Fig. 1e, blue arrows). This suggests that tau^E14^ expression is sufficient not only to induce misfolding and aggregation of tau within the axon, but also to spread as a seed to neighbouring cells and potentially recruit endogenous tau. To more precisely investigate the propagation between neurons we co-cultured tau^E14^ expressing ‘donor’ cells and tau^WT^ ‘acceptor’ cells within a microfluidic device that allows co-culture of two spatially distinct neuronal populations. These two populations can be manipulated independently but are in contact via projecting axons^22,28^. This enabled us to separate tau^E14^ and tau^WT^ expressing neurons and identify individual connecting cells (Fig. 2a-c). We used fluorescence microscopy to analyse the time course of aggregate formation and propagation *in vitro*. Aggregates were first detected at DIV8 in the axons of tau^E14^ expressing donor neurons (Fig. 2e), and the number of axons positive for tau aggregation steadily increased at a rate of 4.9 ± 0.6% per day, until a plateau was reached at DIV18 with 75.5 ± 6.4% of axons containing visible tau aggregates (Fig. 2e). Next, we analysed the percentage of aggregate positive acceptor neurons to assess if propagation occurs in our system. We first detected aggregates in tau^WT^ acceptor axons at DIV10, and their percentage increased over time when connected to tau^E14^ expressing donor neurons (Fig. 2d, e), while no aggregation was detected when tau^WT^ acceptor neurons were connected to GFP expressing control cells (p<0.0001)(Fig. 2d, e). There was an increase of 4.9 ± 0.4% of additional distal axons positive for tau aggregation per day, following the donor neurons with a delay of ~3.7 days, until 77.5 ± 7.8% of distal axons contained tau aggregates, reaching the same plateau as donor distal axons (p>0.999 at DIV24 and DIV26). This demonstrates that within a minimal neuronal circuit free from other cell types, transfer of misfolded tau seeds between primary neurons is rapid and highly efficient. Therefore, spontaneous tau misfolding robustly occurs upon tau^E14^ expression in primary hippocampal neurons, and rapidly and efficiently propagates to connected cells.

**Figure 2.**
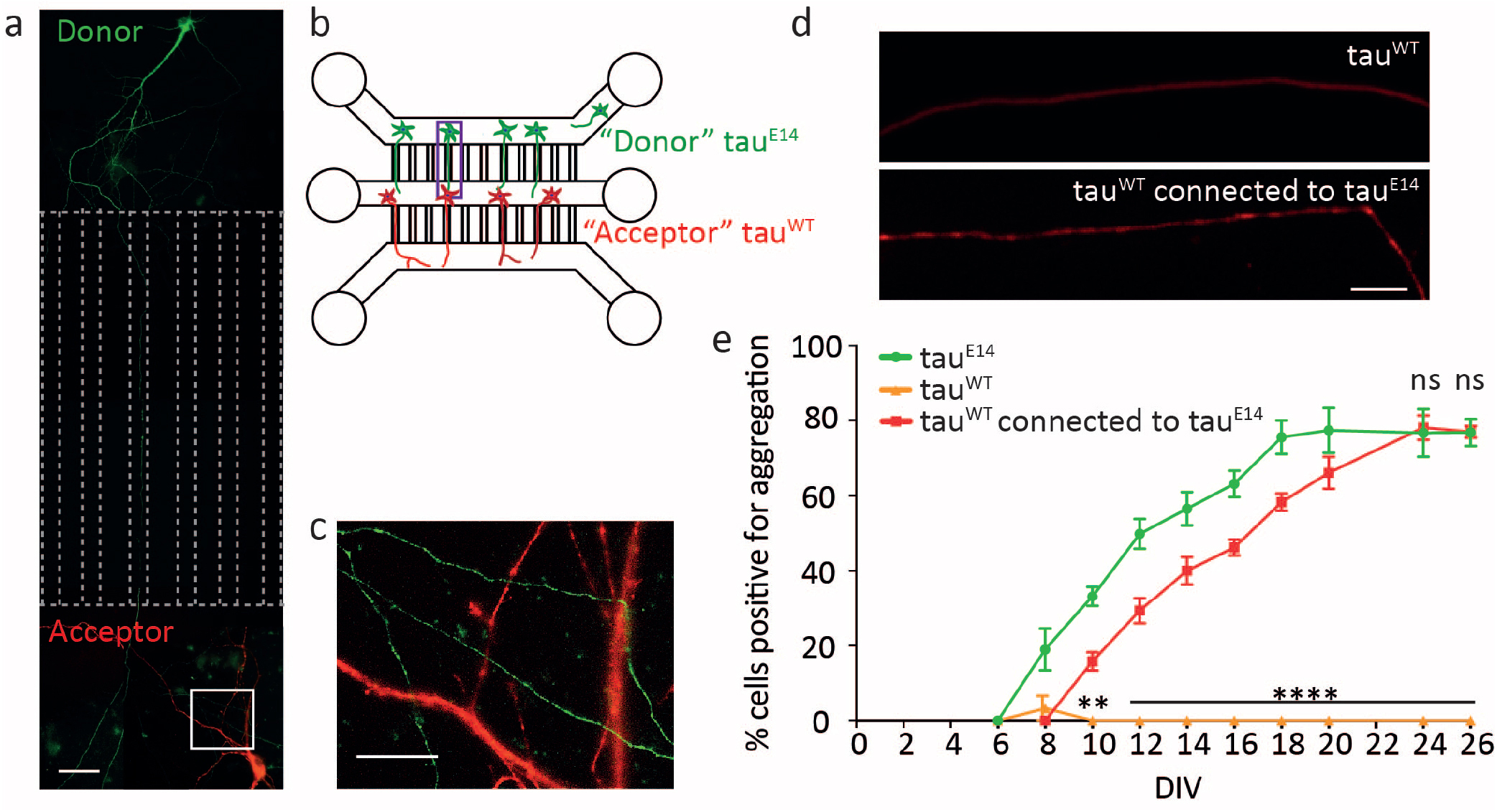
Tau misfolding efficiently propagates to connected neurons. (a) Example image of donor and acceptor cells connected within microfluidic devices. Dashed lines indicate channel boundaries. Scale bar, 15 μm. (b) Schematic of microfluidic device to investigate propagation from donor (tau^E14^, green) to acceptor (tau^WT^, red) neurons. (c) Higher magnification of white box in (a) showing intersections of donor axons and acceptor dendrites. Scale bar, 5 μm. (d) Tau^WT^ expressing acceptor neurons form aggregates when connected to tau^E14^ expressing donor neurons (e) Neurons were transfected at DIV1 and analysed for aggregate formation. Time course of aggregate formation in axons of tau^E14^ expressing donor neurons (green), connected tau^WT^ expressing acceptor neurons (red), and tau^WT^ expressing neurons connected to GFP expressing control cells (orange). N≥12 axons per experiment, 3 independent experiments per time point. Two-way ANOVA with Tukey’s test, F_2,60_=851, ns p>0.999, ** p=0.0028, **** p<0.0001. Error bars=SEM.

### Distinct neuronal subcompartments display a differential vulnerability to tau misfolding and aggregation

In both the donor (not shown) and acceptor neurons, tau aggregates first appeared in the distal axon, and were later detected in the somatodendritic compartment (Fig. 3a). For distal axonal misfolding and aggregation to occur in the acceptor neuron within the oriented setup of the microfluidic chamber (Fig. 2a,b), seeds that have been internalised at the somatodendritic compartment must have first trafficked or propagated to the distal axon. Indeed, we detected GFP-tau^E14^ positive accumulations in the somata of RFP-tau^WT^ expressing neurons (Fig. 3b,d, and movie S1), while no aggregates were visible within the somata of tau^WT^ expressing neurons connected to GFP expressing cells (Fig. 3c,e). The aggregates within the tau^WT^ expressing somata were dual positive for GFP-tau^E14^ and RFP-tau^WT^, demonstrating that the pseudohyperphosphorylated tau had recruited native tau into aggregates (Fig. 3b,d) and suggesting a prion-like propagation. No GFP-tau^E14^ was detected in aggregates of RFP-tau^WT^ expressing axons (not shown), thus further substantiating that tau^E14^ seeded the misfolding of tau^WT^. We confirmed that the misfolding of tau was propagated to the distal axons of acceptor cells using MC1 staining (Fig. 3f). This demonstrates that as well as being sufficient for tau aggregate formation, pseudohyperphosphorylation of tau is sufficient for the prion-like propagation of the conformational change to surrounding neurons. Interestingly, we never detected tau aggregation at the proximal axonal segment of either donor or acceptor neurons, and this axonal subcompartment also remained negative for MC1 staining (Fig. 3f). Together with the observation that visible tau aggregates were first observed in the distal axon of acceptor cells, despite receiving the seeds at the somatodendritic compartment, this suggests that (i) smaller seeds distribute throughout the neuron prior to the formation of visible aggregates, (ii) that tau in the distal axon is most susceptible to aggregation and (iii) misfolded tau does not accumulate in, or is rapidly cleared from, the proximal axonal region.

**Figure 3.**
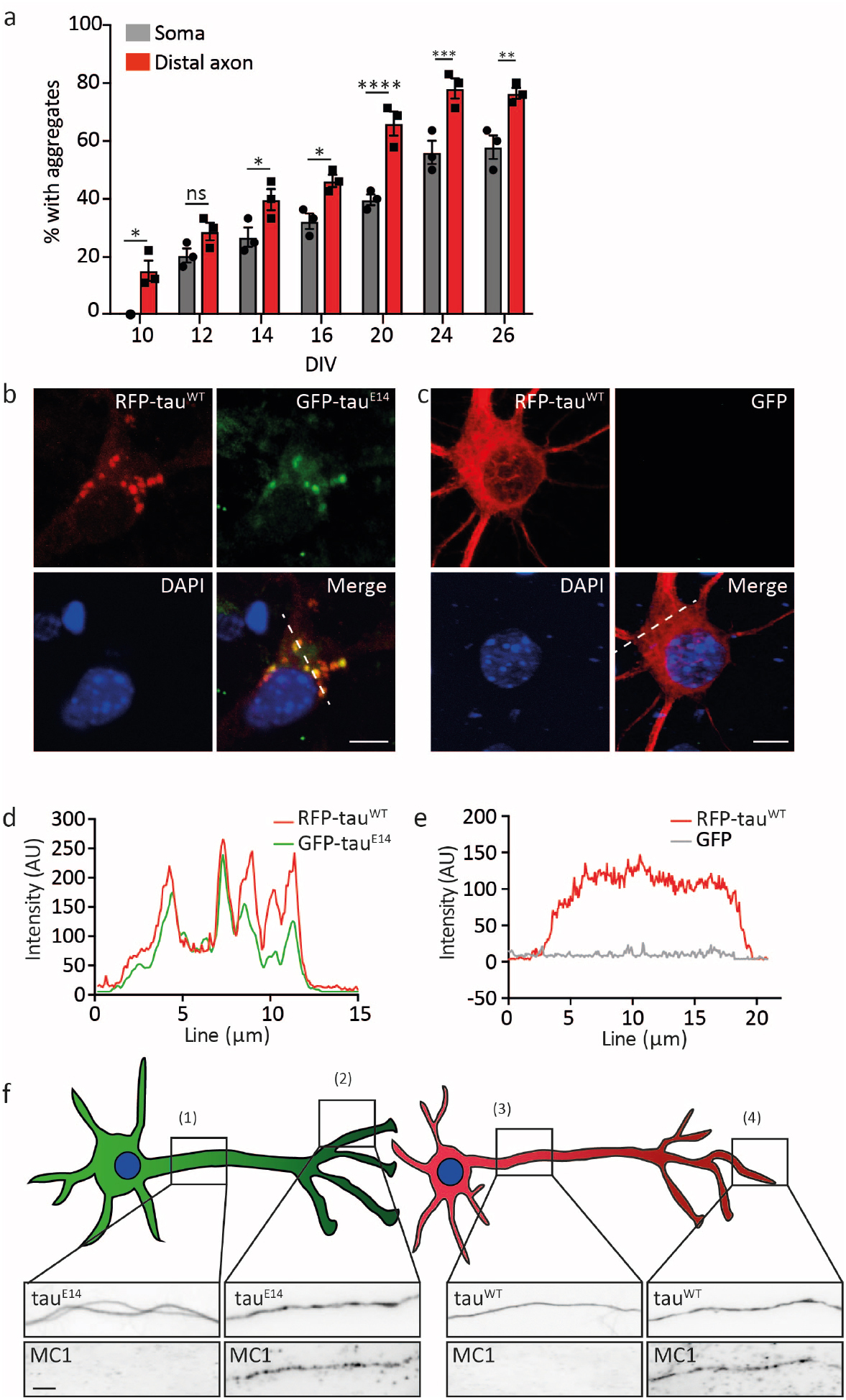
Differential subcompartment vulnerability to tau misfolding. (a) Acceptor somata (grey) and distal axons (red) containing tau aggregates at different time points. Each data point is the mean of one experiment, n≥8 cells per time point per experiment. Two-way ANOVA with Tukey’s test, F_1,4_=68.02, ns p>0.05, * p<0.05, ** p<0.01, *** p<0.001, **** p<0.0001. Error bars=SEM. (b,d) Confocal microscopy shows that GFP-tau^E14^ aggregates are detected in the somata of RFP-tau^WT^ expressing acceptor neurons and overlap with RFP-tau^WT^ aggregates (shown as Z-projection, see also movie S1). Scale bar, 10 μm, imaged at DIV14. (c,e) RFP-tau^WT^ expressing neurons connected to GFP expressing cells do not develop aggregates. Shown as Z-projection. Scale bar, 10 μm, imaged at DIV14. (f) Proximal (1,3) and distal (2,4) axonal segments of tau^E14^ and connected tau^WT^ expressing neurons were counterstained for MC1 and imaged at DIV14. Scale bar, 5 μm.

### Intracellular tau aggregates are not acutely toxic

*In vivo* models show that progression of tau pathology is accompanied by significant neuronal death^4^, and indeed advancement of AD is associated with brain atrophy caused by extensive neuronal loss^1^. Furthermore, exogenous addition of tau is toxic to neurons *in vitro*^17,18,29^, suggesting that tau pathology is associated with neuronal death. However, the direct functional consequence of tau misfolding and aggregation on individual neurons has not yet been identified. Here, we monitored the formation and propagation of tau pathology at single cell level, and thus were able to analyse the state of tau^E14^ expressing cells at the time of misfolding and active tau pathology transmission. Tau^E14^ expressing donor neurons at DIV14 appear intact and undistinguishable from surrounding untransfected cells, as judged by differential interference contrast (DIC) imaging, exhibiting a smooth and intact plasma membrane and healthy nuclear morphology (Fig. 4a). Loss of synaptic connections has been identified as one of the earliest cellular changes in Alzheimer’s disease^30,31^. As tau misfolding and aggregation first occur in the distal axon we visualised a presynaptic marker, synapsin, to assess the density of presynaptic sites. We did not observe any overt differences in the number of presynaptic sites between tau^E14^ expressing neurons and their untransfected surrounding axons (Fig. 4b). This suggests that misfolded tau and tau aggregates are not acutely toxic to hippocampal neurons. Indeed, we did not observe disintegration or death of these neurons across the observed time course, to DIV26. Even at DIV42, five weeks after the first detected propagation of tau pathology to connected neurons (Fig. 2e), the neurons remained intact as judged by DIC imaging and nuclear morphology (Fig. 4c), further dissociating the connection between tau aggregation and acute toxicity. However, at this stage there was a visible decrease in synapsin-positive presynaptic sites along the tau^E14^ expressing axon compared to its surrounding axons (Fig. 4d), suggesting that over time, misfolded tau does lead to presynaptic loss *in vitro*, in line with observations from mouse models and post-mortem brain^30–32^.

**Figure 4.**
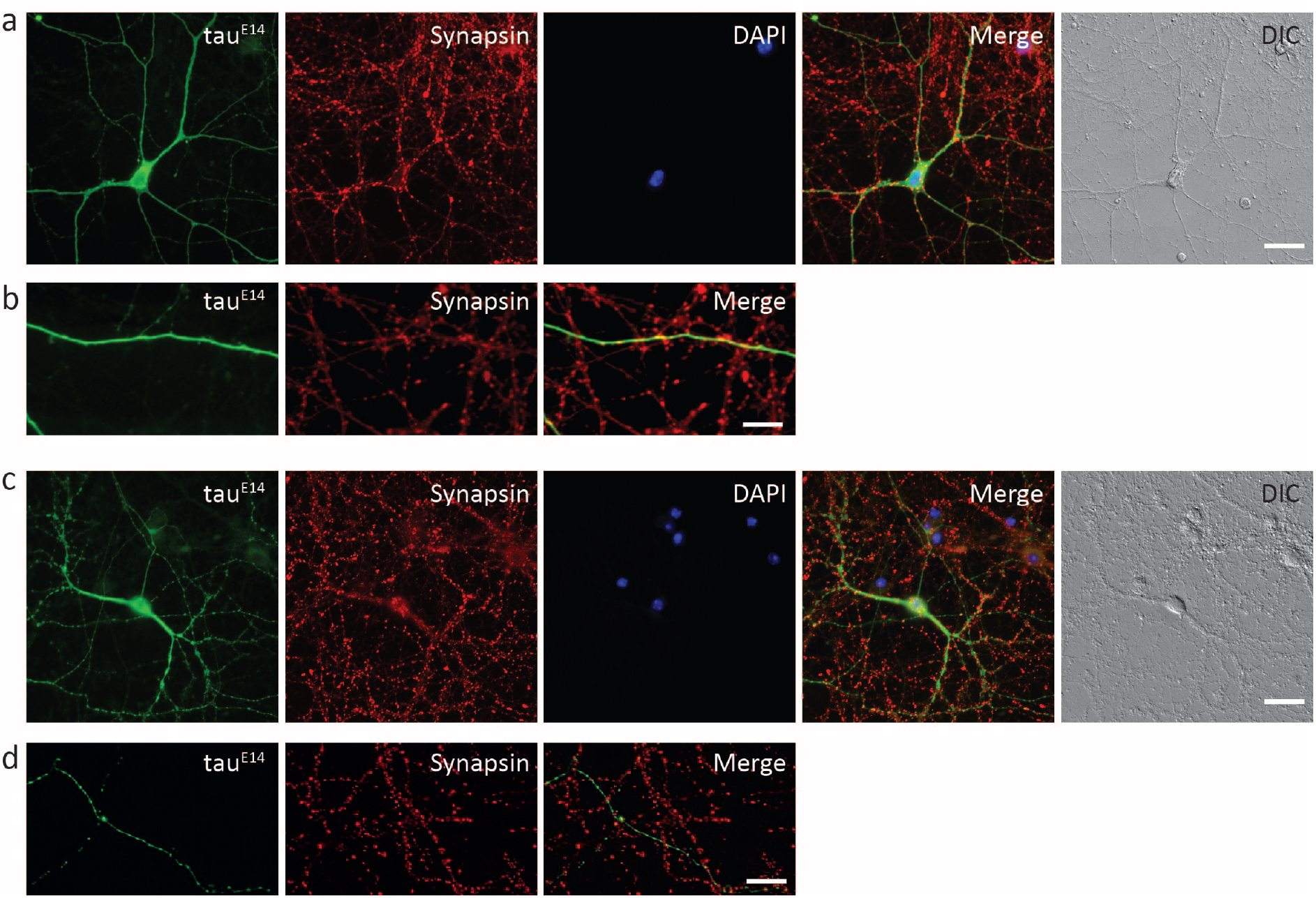
Neurons transmitting tau pathology remain intact. (a,b) A DIV14 tau^E14^ expressing neuron. (a) Overview of neuron stained with synapsin, nucleus stained with Hoechst 33342, and the DIC image showing an intact membrane at the cell body and along the axon. Scale bar, 30 μm. (b) The distal axons of tau^E14^ expressing neurons counterstained with synapsin. Scale bar, 10 μm. (c,d) A DIV42 tau^E14^ expressing neuron. (d) Overview of a neuron stained with synapsin, nucleus stained with Hoechst 33342, and the DIC image showing an intact membrane at the cell body and along the axon. Scale bar, 20 μm. (d) The distal axon counterstained with synapsin. Scale bar, 10 μm.

We next investigated the physiological state of aggregate-containing neurons in more detail to assess functional consequences. Because tau acts in microtubule stabilisation^33^, and its dysfunction leads to axonal transport deficits^34,35^, we first assessed lysosome dynamics. Lysosomes are transported bi-directionally along axons^36^. At DIV14 there was a significant decrease in the number of lysosomes present at the distal axon of tau^E14^ expressing neurons compared to tau^WT^ controls (Fig. 5a,b). Of those lysosomes present, a lower percentage were moving within tau^E14^ axons (21.1 ± 3.9%) compared to tau^WT^ controls (45.8 ± 5.8%)(Fig. 5c-e; see also movies S2, S3), confirming an early axonal transport dysfunction in the presence of tau misfolding.

**Figure 5.**
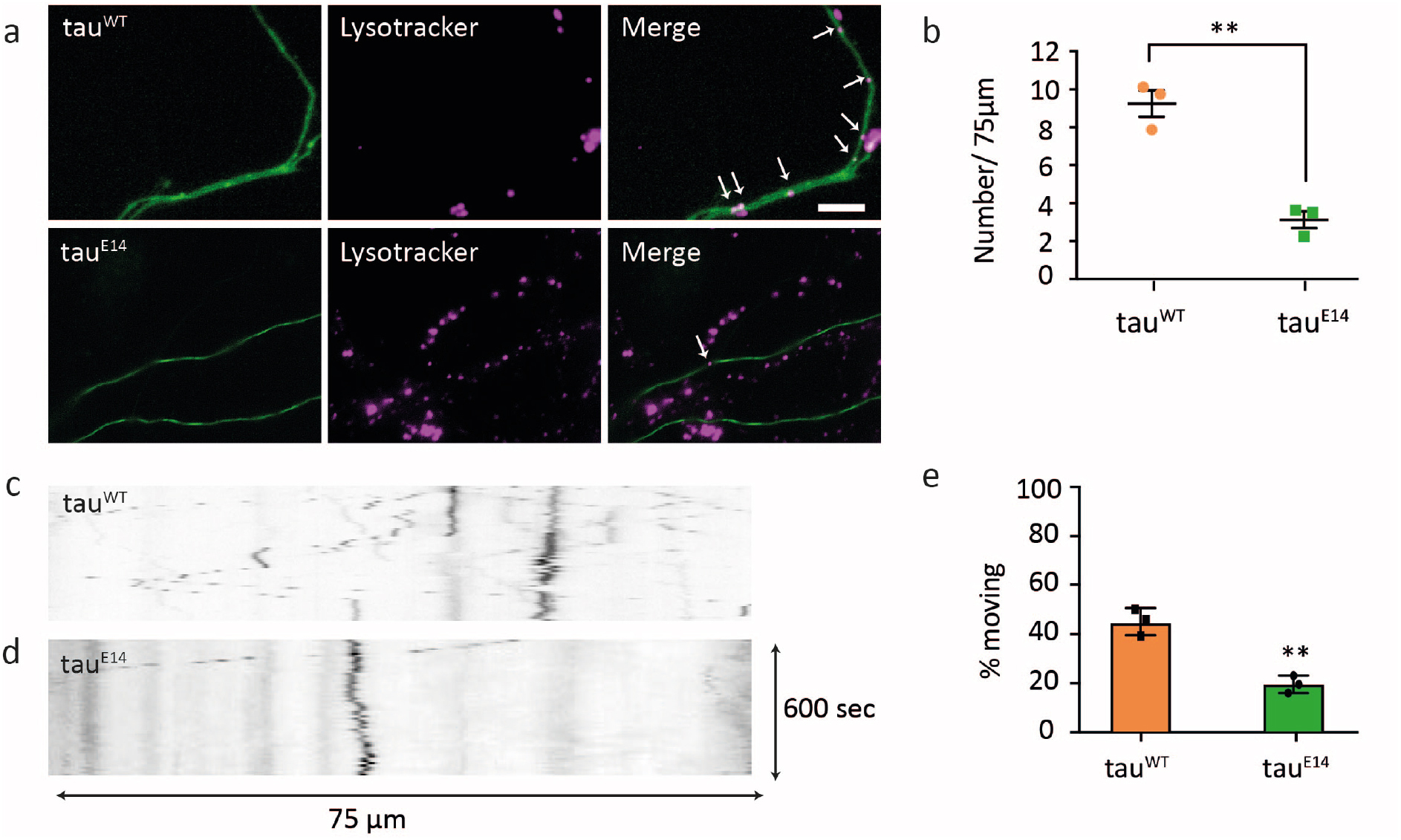
Tau^E14^ expressing neurons show selective axonal defects at DIV14. (a-e) Imaging of lysosomes at the distal axon reveals (a,b) a reduction in the number of lysosomes (indicated by white arrows) in tau^E14^ expressing distal axons compared to tau^WT^ expressing cells. N=8 cells per experiment, each point is the mean of one experiment. Unpaired t-test, df=4, t=7.444, p=0.0017 (c-e) Analysis of lysosome dynamics in tau^E14^ and tau^WT^ expressing axons. (c,d) Representative kymographs of tau^WT^ and tau^E14^ expressing axons, see also movies S2, S3. (e) Transport analysis of lysosomes. Each data point is the mean of one experiment, n≥8 cells per experiment. Error bars=SEM. Unpaired t-test, df=4, t=6.143, p=0.0036.

To confirm our observation that neurons expressing tau^E14^ remained viable in the presence of misfolded tau we then visualised spontaneous activity using calcium imaging (Fig. 6a,b and movies S4, S5). Both tau^WT^ and tau^E14^ expressing neurons displayed calcium fluxes that were sensitive to tetrodotoxin, showing that they were driven by voltage-gated sodium channels. This demonstrates that energy-dependent processes were intact, and suggests that neurons remain electrically competent in the presence of misfolded and aggregated tau. To investigate the electrical competence in more detail, we performed an electrophysiological analysis on the cells with whole-cell patch clamp. When current clamped, both tau^E14^ and tau^WT^ expressing neurons were capable of responding to positive current injections with action potentials of a similar amplitude (Fig. 6c). There were no significant differences in the minimum current required to evoke a single action potential (rheobase); with tau^E14^ and tau^WT^ expressing neurons requiring 77 ± 40 pA and 83 ± 25 pA respectively (p = 0.53)(Fig. 6d). We further found no significant differences in the input resistance between tau^E14^ (694 ± 367 MΩ) and tau^WT^ (578 ± 101 MΩ) expressing cells, (p = 0.64)(Fig. 6e), and the resting membrane potentials of tau^E14^ (−73.8 ± 2.4 mV) and tau^WT^ (−73.3 ± 3.5 mV) expressing neurons do not differ, (p = 0.58)(Fig. 6f). This shows that despite possessing and transmitting tau pathology, donor neurons retain their ability to maintain a normal resting membrane potential and fire action potentials to the same extent as tau^WT^ control neurons.

**Figure 6.**
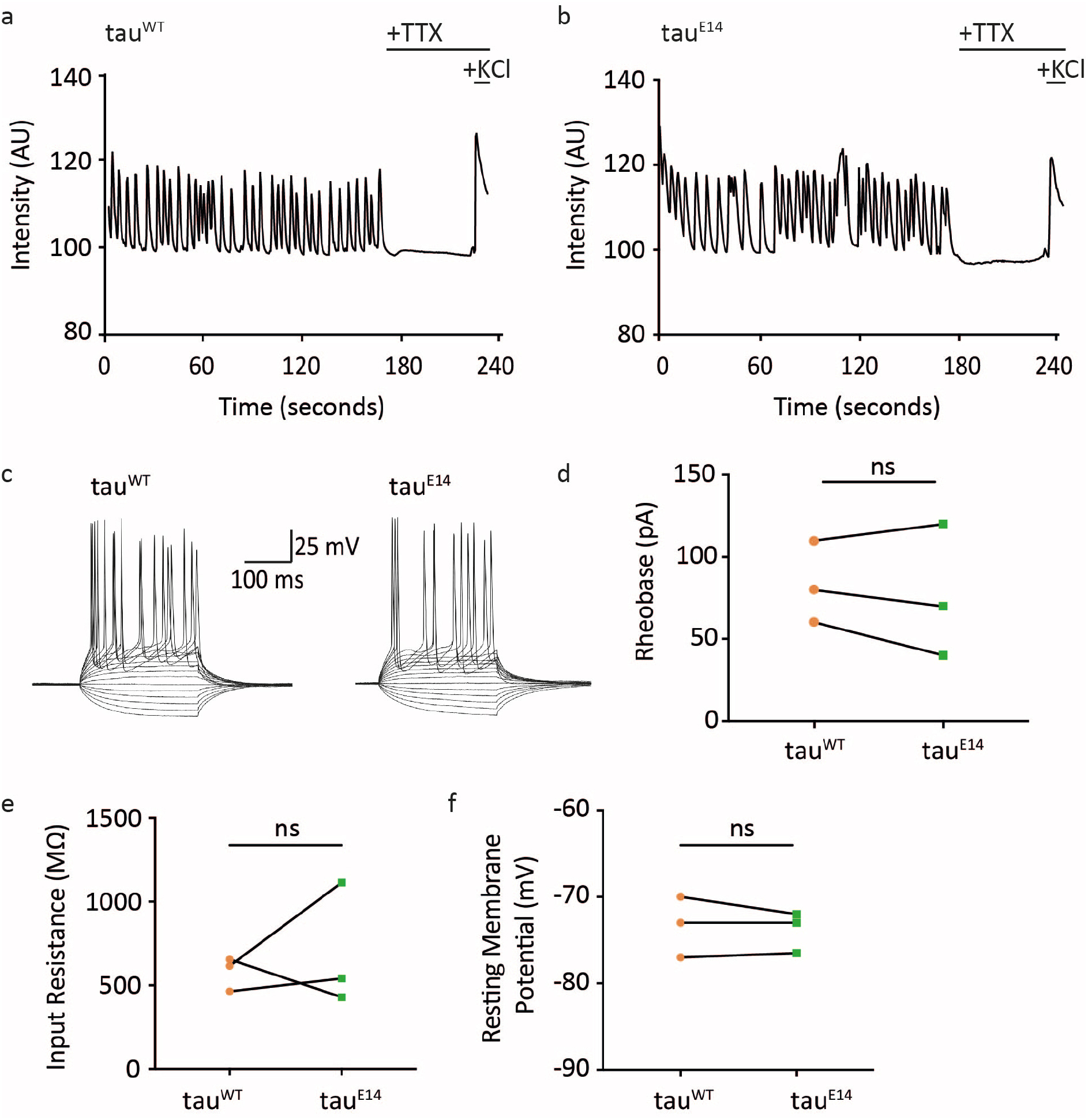
Tau^E14^ neurons are electrically competent. (a,b) Basal calcium activity of a DIV14 (a) tau^WT^ (b) and tau^E14^ expressing neuron, which is silenced on application of 1 μM TTX. Addition of 100 mM KCl confirms viability at the end of acquisition. See also movies S4, S5. (c-f) Electrophysiology on DIV14 neurons. (c) Sample traces of current injection into patch clamped neurons. (d) Rheobase of tau^WT^ and tau^E14^ expressing neurons show a similar minimal current is required for action potential initiation. Paired t-test, df=2, t=0.7559, p=0.5286. (e) Input resistance of tau^WT^ and tau^E14^ expressing neurons shows no significant difference. Paired t-test, df=2, t-0,5542, p=0.6363. (f) Resting membrane potentials recorded for tau^WT^ and tau^E14^ expressing neurons. Paired t-test, df=2, t=0.6547, p=0.5799. For (d-f), N≥3 cells per experiment, each point = median of 1 experiment.

Together, these data demonstrate that phosphomimetic tau misfolds and aggregates in the absence of exogenous seeds, and that transmission of tau misfolding to healthy neurons is an active and efficient process that is accompanied by a selective neuronal dysfunction, but the presence of misfolded and aggregated tau does not compromise neuronal excitability and is compatible with longer-term neuronal viability.

## Discussion

Using a minimalistic *in vitro* model of connected neurons, we have shown that phosphomimetic tau misfolds in the absence of exogenous seeds and forms tau aggregates. This tau pathology is robustly propagated from the neuron expressing the phosphomimetic protein to surrounding neurons. We further demonstrate that neurons containing tau aggregates display no neurodegeneration. Instead, they remain intact and retain electrical competence at the time of tau transmission. While the presence of tau aggregates leads to a selective impairment in axonal transport, this does not impact on neuronal viability. These observations show that tau misfolding and aggregation do not induce acute toxicity in hippocampal neurons.

Previous studies examining the mechanisms of tau propagation have added varying exogenous species of tau seeds to induce misfolding^5,11,22^. We here show that the negative charges conferred by a phosphomimetic tau that simulates hyperphosphorylation at 14 disease-related sites are sufficient to initiate misfolding within a living neuron, in absence of exogenous seeds. This misfolded tau forms aggregates and propagates tau seeds from presynaptic to postsynaptic neurons. AD is associated with an imbalance of kinase and phosphatase activities^37^ that leads to aberrant hyperphosphorylation of tau, and these hyperphosphorylated species are detected within NFTs^38^. Our observation implies that a post-translational modification such as phosphorylation in isolation is sufficient to initiate the cascade of tau misfolding, aggregation and propagation.

The compartmented setup of our propagation assay allowed the monitoring of individual neurons that formed or received misfolded tau species. This showed unexpectedly fast and robust tau propagation from neuron to neuron *in vitro*, with a near complete transmission efficiency. This is at odds with findings that show Braak staging progresses in patients over a matter of years to decades^39^. Our study used pure murine hippocampal neurons, cultured in the absence of other cell types. The efficiency of transmission may be due to a lack of a glial population, which display tau pathology in patients with tauopathy^40–42^, as well as in *in vivo*^43^ and *in vitro*^44^ models of tauopathy. Therefore, glial populations may play a role in clearance of secreted pathogenic tau, and their absence in our setup revealed an under-appreciated intrinsic high efficiency of tau release and re-uptake in neurons. This high efficiency of neuron-to-neuron tau propagation within our connected system further suggests that a physiological process of protein transmission may be at play that is occurring in healthy neurons, but revealed under pathological conditions through transmission of a conformationally altered species. This idea is reinforced by the observation that healthy tau is secreted from intact neurons in an activity-dependent manner^45^.

Our minimalistic setup allows direct investigation of neurons that initiate misfolding or receive transmitted tau seeds at single cell and subcellular level. This revealed a compartmentalisation of tau aggregation within individual neurons. We observed that tau aggregate formation within an individual neuron begins within the distal axon, regardless of whether misfolding was originally initiated within the cell, or transmitted to it. This spatial organisation holds true despite our oriented setup of connected neurons, where tau seeds first enter the receiving neurons at the somatodendritic region; the site furthest from the distal axon where aggregation is initiated. Only later were aggregates visible in the somata. Furthermore, we did not observe tau aggregates in the proximal axonal segment, even at stages where clear aggregates were visible in both the soma and distal axon. This suggests either a selective vulnerability of the distal axonal segment, or protection against tau misfolding in proximal axonal regions. The region of the axon initial segment has been shown to act as a barrier for select isoforms of tau proteins, potentially due to increased microtubule dynamics in this area^46,47^, which may play a role in the lack of retention of misfolded tau within this region. However, it is clear that misfolded tau is able to cross this barrier and cause distal axon pathology.

Our ability to monitor tau misfolding within single cells allowed us to directly assess the physiological effects that tau pathology has on neurons. We found that despite containing misfolded tau and tau aggregates for prolonged periods of time, neurons remained alive, intact, and retained energy dependent processes, displaying a selective dysfunction in axonal transport. Previous studies showed that acutely added tau seeds can induce toxicity and cell death via calcium dysregulation^17,18,29^. However, we found that neurons containing tau aggregates had intact calcium fluxes, physiological resting membrane potentials, and were able to elicit action potentials. Furthermore, we also showed cell survival for extended periods of time after tau aggregate formation, as we observed neuronal viability for weeks after initial misfolding and propagation. Our data therefore shows that misfolded tau and tau aggregates are not immediately toxic to neurons. Instead, they initiate a selective axonal transport dysfunction that precedes synaptic loss, and in isolation does not result in axonal degeneration or cell death.

Furthermore, we provide the first direct evidence for the suggestion that disintegration or death of neurons containing tau seeds does not need to occur in order to propagate tau pathology^20^, and show that tau propagation precedes synaptic or neuronal degeneration. This is in agreement with a recent study showing that spread of tau seeds precedes tangle pathology in the human brain^21^. In addition, prion seeds are detectable throughout all brain regions in mice infected with prion disease, while degeneration is region specific^48^. Together, these data and our findings suggest that the idea that presence of misfolded protein in isolation does not determine neurodegeneration^48^ may hold true across different protein misfolding diseases. This information, combined with a potential physiological transmission of tau between neurons, raises the possibility that at the stage of diagnosis, tau seeds may have spread throughout wider brain regions, having passed the time point where a sequestration of free tau seeds in isolation is effective in halting disease progression.

However, our data imply that because the affected neurons are intact and viable at the early stages of tau misfolding, and because neither hyperphosphorylated tau nor tau aggregates are immediately toxic to neurons, neurons with tau pathology could potentially be rescued with a therapeutic disease modifier.

## Online methods

### Plasmids

The following plasmids were used: pRFP-N1, pEGFP-C3 (Clontech), pRK5-EGFP-tau^WT^ and pRK5-EGFP-tau^E14^ were a gift from Karen Ashe^25^ (Addgene plasmids #46904 and #46907), GCaMP6 was a gift from Douglas Kim^49^ (Addgene plasmid #40753), R-GECO was a gift from Robert Campbell^50^ (Addgene plasmid #45494). RFP-tau^WT^ and RFP-tau^E14^ were created by excising the GFP fragment of pRK5-EGFP-Tau^WT^ at Cla1 and BamH1 sites, and replacing it with RFP, which was amplified by PCR with forward primer 5′-CATGATCGATATGGCCTCCTCC-3′ and reverse primer 5′-CATGGGATCCGGCGCCGGT-3′.

### Cell culture and transfection

All experiments were carried out in accordance with the Animals (Scientific Procedures) Act 1986 set out by the UK Home Office. Primary cultures were prepared as described previously^51^. Briefly, hippocampal neurons were isolated from embryonic day 15-18 C57BL/6 mouse brains. Dissociated neurons were plated in Neurobasal medium supplemented with 2% B27 and 0.5 mM GlutaMAX (Gibco) at a density of 7000 cells/μl in microfluidic devices, and 150,000 cells/ml in glass bottom dishes. Partial medium changes were performed on the devices every 2-3 days. On DIV1, neurons were transfected using Lipofectamine 2000 as described previously^51^.

### Microfluidic devices

Custom microfluidic devices were manufactured based on existing designs^52,53^ (see supplemental file 2). Devices were replicated using polydimethylsiloxane (PDMS, Sylgard 184, Dow Corning) with a 10:1 ratio of elastomer to curing agent. PDMS mix was cured at 60 °C for at least 90 minutes, and inlets were punched using a 5-mm-diameter biopsy punch (Miltex). Devices were washed in 70% EtOH for one hour and dried before use. Devices were mounted onto 22×55 mm coverslips (Smith Scientific), which had been pretreated with 0.1 mg/ml poly-*D*-lysine (Sigma). Device channels were filled with supplemented Neurobasal medium and incubated overnight before addition of cells.

### Immunocytochemistry

Neurons were fixed in 4% paraformaldehyde in PBS for 10–15 minutes, washed with 50 mM ammonium chloride in TBS for 5 minutes, and permeabilised in 0.1% Triton-X100 in TBS for 5 minutes at room temperature (RT). The cells were then blocked for 30 minutes in 10% goat serum in TBS at RT. Cells were incubated for 1 hour at RT or overnight at 4°C in the following primary antibodies: MC1 (1:300, gift from Peter Davies^27^), synapsin-1 (D12G5, 1:1000, Cell Signalling), ankyrin G (N106/36, 1:300, NeuroMab). Primary antibody was washed off in TBS, and Hoechst (33342) was added to the second wash to stain nuclei (1:3000, Thermo Fisher). This was followed by incubation for 30 minutes at RT in Alexa fluorescent-conjugated secondary antibodies: anti-mouse Alexa647 or anti-rabbit Alexa555 (Invitrogen). Coverslips were mounted onto microscope slides using Mowiol (Sigma).

### Microscopy

Fixed cell images for axonal length analysis were taken on a Zeiss Axioplan Fluorescence Microscope equipped with a HBO103 Mercury lamp for illumination, a Qimaging Retiga 3000 monochrome CCD camera (Photometrics, UK), 20x/0.4NA and 40x/0.75NA Plan-Neofluar objectives, using Micro-manager software (Vale lab, USA). Fluorescent and differential interference contrast (DIC) images of cells in devices were obtained using a 60x/1.42NA Oil Plan APO objective on a DeltaVision Elite system (GE Life Sciences) with SSI 7-band LED for illumination and a monochrome sCMOS camera, using SoftWoRks software (version 6). Confocal images were taken on a Leica SP8 laser scanning confocal microscope using a 63x/1.30NA HC Pl Apo CS2 glycerol immersion objective, with a PCO Edge 5.5 sCMOS camera. Lasers used for illumination were continuous wave solid state lasers at 405 and 561 nm, and a continuous wave argon gas laser at 488 nm. Live cell imaging was performed on the DeltaVision Elite system (GE Life Sciences). For imaging of lysosomes, DIV14 neurons were incubated with 25 nM LysoTracker Deep Red (Thermo Fisher) for 20 minutes at 37°C. LysoTracker solution was then removed and replaced with supplemented NBM containing 50 mM HEPES-NaOH, pH 7.4. Images were taken at 0.2 Hz for 5 minutes using 80 ms exposure with 5% light. For calcium imaging, neurons were cotransfected with GCaMP6 and RFP-Tau^E14^ or RFP-Tau^WT^, or R-GECO and GFP-Tau^E14^ or GFP-Tau^WT^ and imaged at DIV14, using 100 ms exposure and 2% light. Images were taken at 2 Hz for 6 minutes. After 3 min, 1 μM tetrodotoxin (Sigma) was added, followed after 2 minutes by 100 mM KCl.

### Image analysis

Overview images were reconstructed from multiple single images using Autostitch software (University of British Columbia). Images were processed and analysed using ImageJ software (NIH), and its plugins NeuronJ^54^ and Iterative Deconvolution (Bob Dougherty). Axonal length was defined as the longest axonal branch from the longest neurite. Distal axon was defined as a 75 μm stretch of axon at, or near to, the terminal of an axon branch, and proximal axon was defined as a 75 μm long stretch of axon measured beyond the first 50 μm of axon protruding from the cell body, therefore beyond the axonal initial segment (Fig. S2). Intensity profiles along a line were generated using plot profile, and kymographs using the MultipleKymograph plugin on ImageJ. Lysosomes which displaced greater than 50 μm over the course of the time lapse were considered moving and all others stationary. For calcium imaging, a circle was drawn around the soma of the expressing neuron, and analysis was generated using the ‘intensity v time’ stacks function of ImageJ.

Aggregate analysis was performed using Matlab. Briefly, the fluorescence intensity values of paired control and experimental axons were measured. A baseline of each individual axon is set by subtracting the 10^th^ percentile from all values, and any point lying outside 5 times the mean of the deviations of all control axons from the experiment is scored positive, i.e. an aggregate containing point. The percentage axonal stretch scoring positive is determined, and any axon with >10% of its length outside the allowed deviation is scored as aggregate containing (Fig S1).

### Electrophysiology

Cells were cultured on coverslips and transfected with RFP-Tau^WT^ or RFP-Tau^E14^. For patch clamp recording cells were perfused with oxygenated (95% O_2_, 5% CO_2_) artificial cerebrospinal fluid (CSF) which contained (in mM): 126 NaCl, 3 KCl, 1.25 NaH_2_PO_4_, 2 MgSO_4_, 2 CaCl_2_, 26 NaHCO_3_, and 10 glucose, pH 7.3–7.4. at a rate of 1–2 ml/min. Recordings were performed under visual control. Patch pipettes (4–6 MΩ) were pulled from thick-walled borosilicate glass tubing and filled with a solution containing (in mM): 110 potassium-gluconate, 40 HEPES, 2 ATP-Mg, 0.3 GTP, 4 NaCl (pH 7.25 adjusted with KOH; osmolarity 280 mosmol/l). Recordings were carried out at room temperature using an amplifier Axopatch 200B. After measurement of intrinsic membrane potential, if necessary, current was injected to maintain the membrane potential −75 ± 5 mV. All membrane potentials recorded were corrected off-line for liquid junction potential of −10 mV measured directly. Current pulses of increasing amplitude were used to test excitability in current clamp. Input resistance was measured in voltage clamp with 2 mV pulses. Signals were low-pass filtered at 5 kHz and sampled at 20 kHz with 16-bit resolution, using a National Instruments analogue card, and custom software written in Matlab and C (MatDAQ, Hugh Robinson, 1995–2013). All analysis was performed in Matlab.

### Statistics

Statistical analysis was performed using GraphPad Prism 6 (Ver 6.00m Graph Pad Software Inc.). All experiments contain data from a minimum of 3 independent dissections, with an individual experiment defined as the cells derived from embryos of one mouse. All data in text are expressed as mean ± SD. Statistical analyses were performed using a t-test for comparison of two groups or an ANOVA for comparison of 3 or more groups, with details provided in figure legends.

### Data availability

The data that support the findings of this study are presented in the paper and are available from the corresponding author upon reasonable request.

## Acknowledgements

We thank Professor Vincent O’Connor, Dr Amrit Mudher and Aleksandra Pitera for critically reading the manuscript. This work was supported by Alzheimer’s Research UK (grant numbers ARUK-PhD2014-10 and ARUK-PPG2017B-001).

## Author contributions

KD conceptualised the study. KD and GIH designed, performed and analysed the experiments, and wrote the manuscript. JJW and GIH designed and optimised the microfluidic devices. MVC performed patch clamping experiments.

## Supplementary material

### Supplemental file 1

#### Supplemental methods: Aggregate analysis

Aggregate analysis was performed in Matlab (see script below). Plot profiles of 75 μm long axonal stretches were generated from 16-bit images, and the pixel intensity values were analysed (Fig. S1). To assess normal fluorescence fluctuations, a selection of intensity values derived from 50 control tau^WT^ axons across the time course were randomly chosen, and each was zeroed to its 10^th^ percentile. The mean+5 standard deviations of these values was calculated as 500 arbitrary units (a.u.). Therefore, tau^WT^ control axons with values above 500 were excluded from generating experimental control means (script line 4, Fig. S1b), but included in the final analysis.

The fluorescence values for at least 12 control tau^WT^ and experimental tau^E14^ 75 μm long axonal stretches were analysed per timepoint, with three separate experiments per time point. The mean of the standard deviations of the control axons not excluded by the 500 a.u. cut-off was calculated (script line 7). Any individual fluorescence value of tau^WT^ or tau^E14^ axons lying 5 times outside this mean was identified to be an aggregate-containing point (script lines 8 and 16 respectively). The sum of the aggregate-containing points was calculated for control tau^WT^ and experimental tau^E14^ axonal stretches, and from this the percentage of aggregate-containing values along an axonal stretch was calculated (script lines 11 and 19 respectively). Any axonal stretch that contains over 10% of its fluorescence values, i.e. over cumulative 7.5 μm of the analysed length, as aggregate-containing values was identified as axon positive for tau aggregation (script lines 12 and 20, Fig. S1c). This cut-off allows for intensity variations due to e.g. crossing axons to be discounted. Finally, the percentage of cells positive for tau aggregation is calculated for tau^WT^ and tau^E14^ expressing neurons (script lines 12 and 20 respectively, Fig. S1d).

1- controlbase=prctile(control,10);
2- controlzero=control-controlbase;
3- max=max(controlzero);
4- include=max<500; %500 determined as described above
5- controzero2=controlzero(:,include);
6- deviation=std(controlzero2);
7- mean=mean(deviation);
8- logicctl=controlzero>5*mean;
9- sumcontrol=sum(logicctl);
10- [n,x]=size(control);
11- percentagecontrol=sumcontrol/n;
12- finalcontrol=sum(percentagecontrol>0.1)/x*100;
13- expbase=prctile(exp,10);
14- expzero=exp-expbase;
15- logicexp=expzero>5*mean;
16- sumexp=sum(logicexp);
17- [m,y]=size(exp);
18- percentageexp=sumexp/m;
19- finalexp=sum(percentageexp>0.1)/x*100;

**Figure S1.**
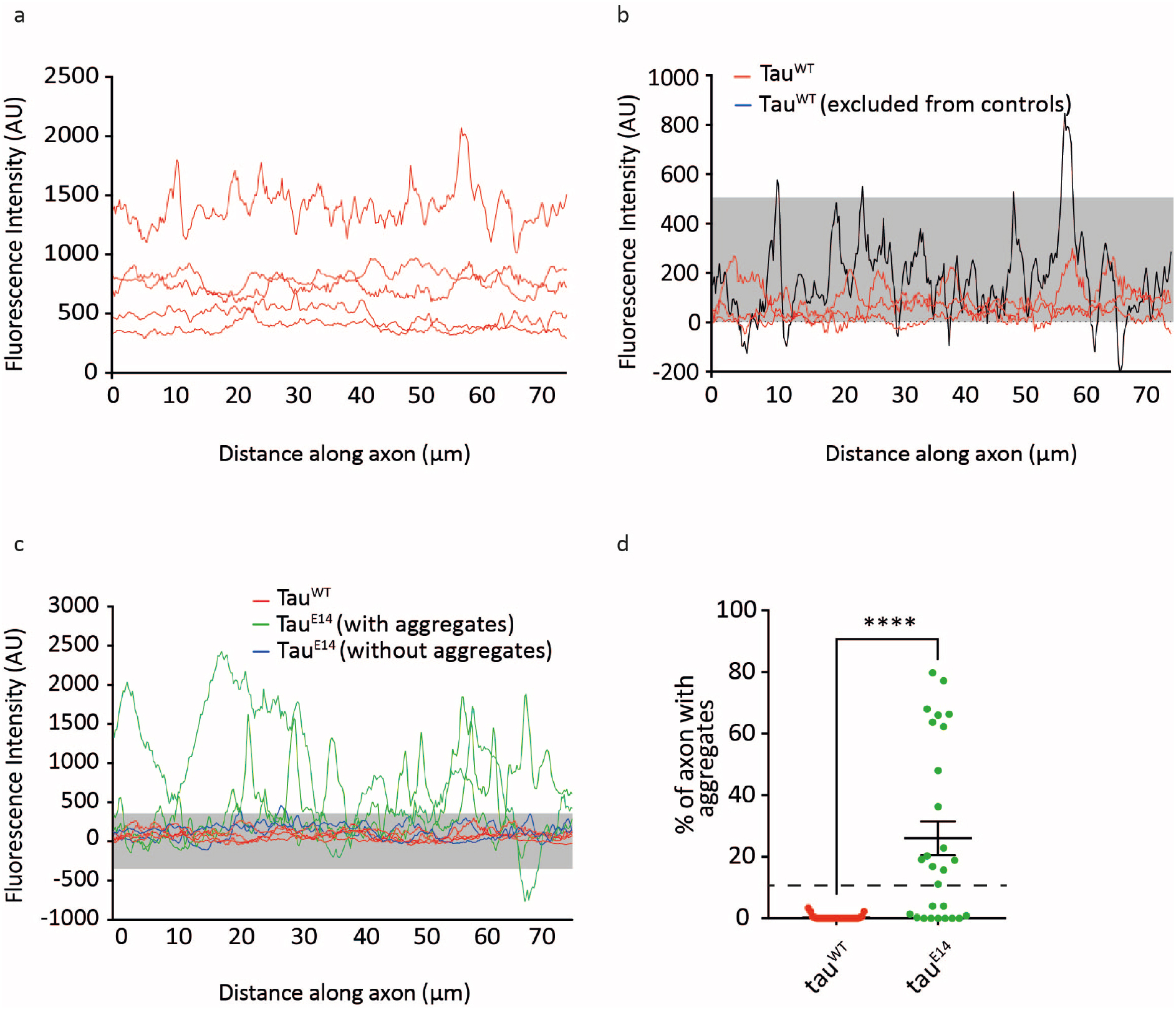
Aggregation analysis. To analyse aggregate formation, (a) intensity profiles from ≥12 tau^WT^ expressing neurons from each time points were generated (see also Fig. 1d). (b) To correct for different expression intensities and offset values to a baseline, the 10^th^ percentile of each trace was calculated and deducted from all values. The mean and standard deviation of intensity values of 30 random axonal stretches were calculated, and any axonal stretch containing values outside this mean+5x standard deviation (grey box, 500 a.u.) was excluded for the purposes of generating a control set but included in the final analysis (black line). (c) From this, we defined the mean+ 5x standard deviation as allowed intensity variation in our final aggregate analysis (c, grey box). (d) Axons with over 10% of their values outside this allowed range were determined as aggregate positive.

**Figure S2.**
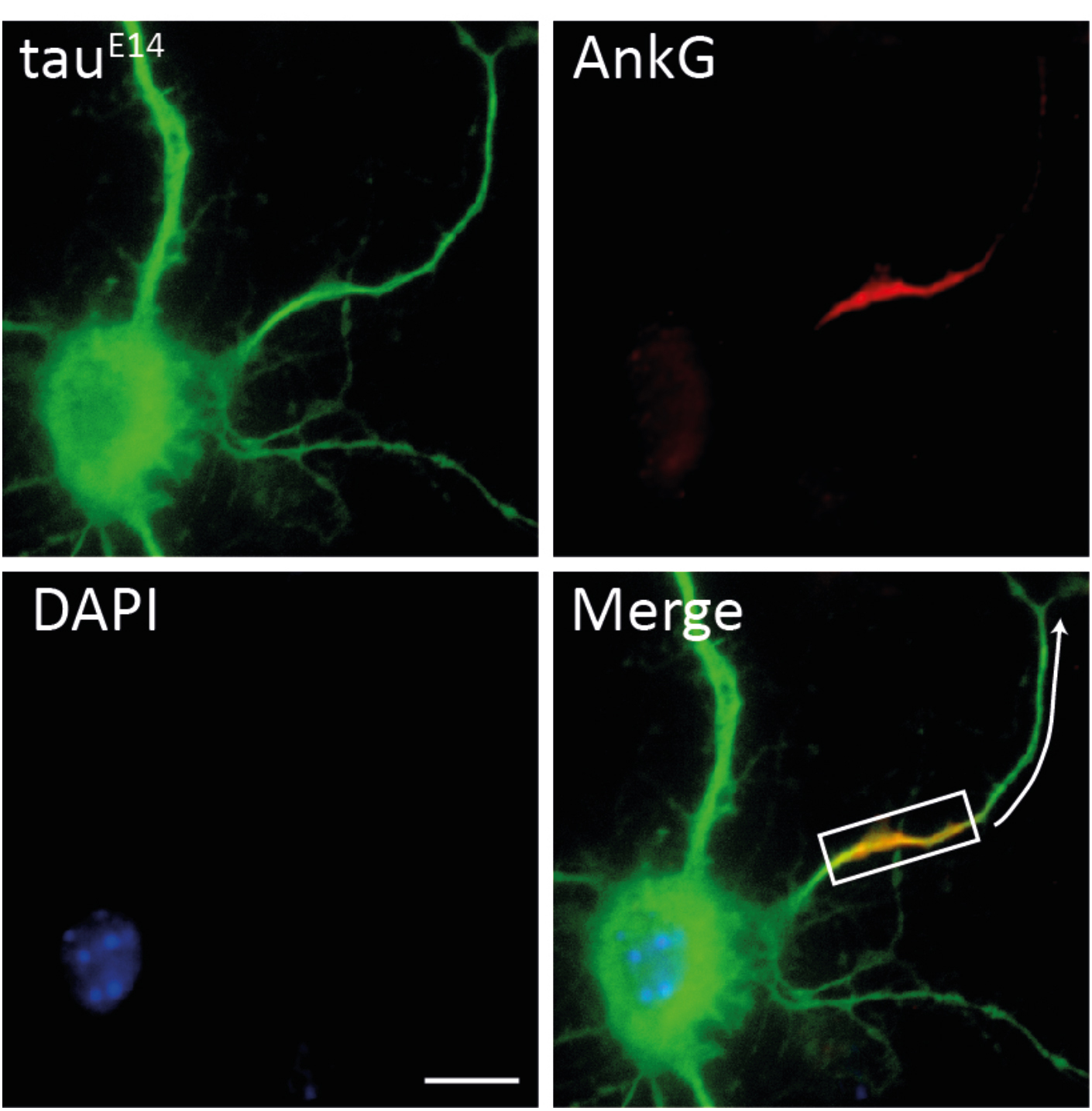
Axon initial segment staining for identification of proximal axon. Axon initial segment identified with AnkyrinG staining. Proximal axon was measured as the first 75 μm axon extending after the axon initial segment (white arrow). Scale bar, 5 μm.

### Supplemental file 2

Images of mask layers for the microfluidic device, provided as DraftSight (.dwg) files.

Layer 1 – height 3 μm.

Layer 2 – height 50 μm.

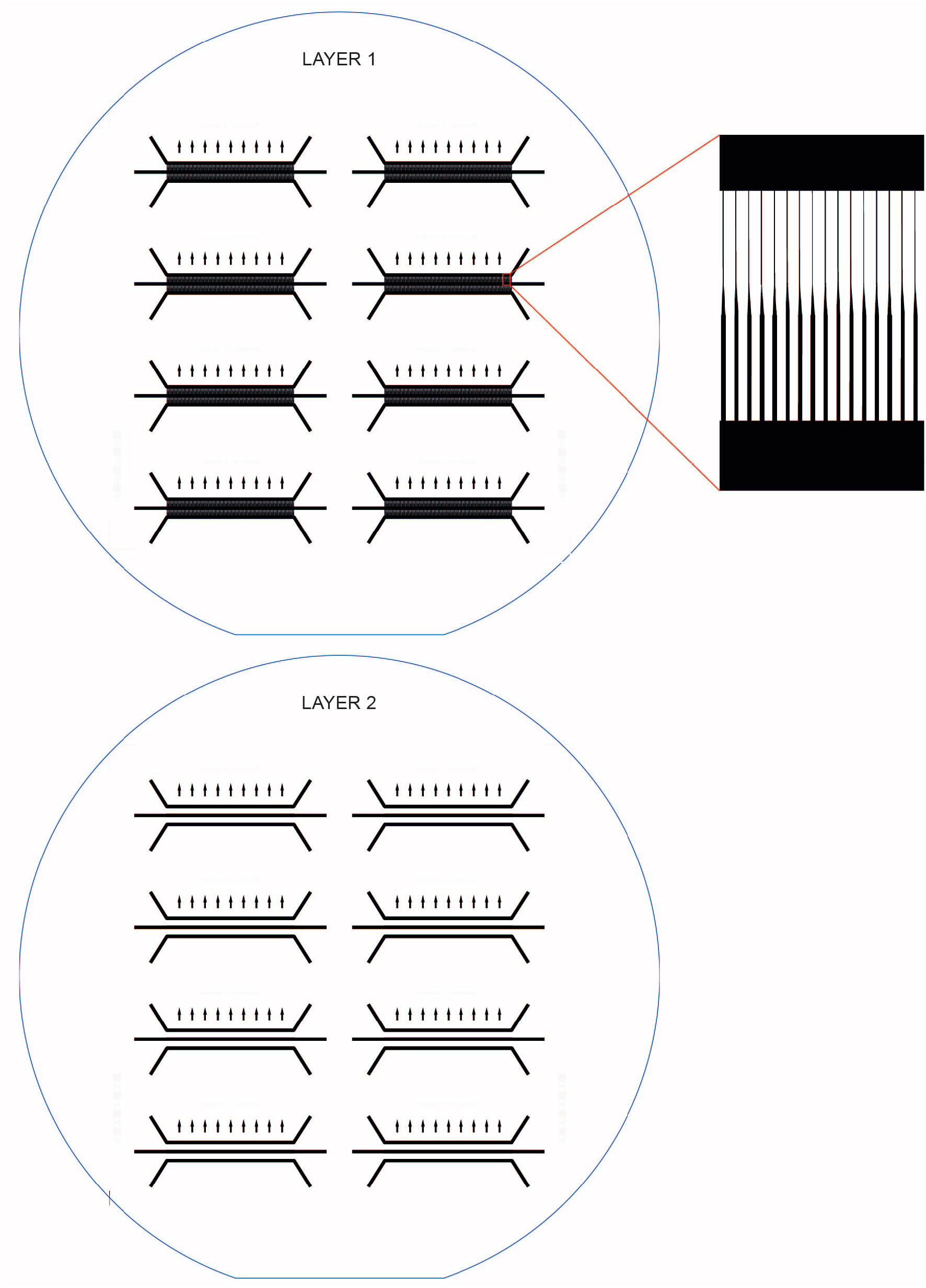

## Supplemental movies

**Movie S1. Tau^E14^ aggregates propagate to acceptor cells and recruit of tau^WT^.** This video shows a rotation of a 3D confocal image of an acceptor soma. This video consists of 200 frames and has an image size of 65.1 x 36.6 μm. See also Fig. 3b.

**Movie S2. Axonal transport of lysosomes in a tau^WT^ expressing neuron.** This video shows the axon of a neuron transfected on DIV1 with tau^WT^, and incubated on DIV14 with LysoTracker Deep Red for 20 minutes prior to imaging. Images were taken at 0.2 Hz. This video consists of 64 frames, and has an image size of 59.2 x 14 μm. See also Fig.5a.

**Movie S3. Axonal transport of lysosomes in a tau^E14^ expressing neuron.** This video shows the axon of a neuron transfected on DIV1 with tau^E14^, and incubated on DIV14 with LysoTracker Deep Red for 20 minutes prior to imaging. Images were taken at 0.2 Hz. This video consists of 64 frames, and has an image size of 63.2 x 8.4 μm. See also Fig.5a.

**Movie S4. Calcium activity in a tau^WT^ expressing neuron.** This video shows a neuron cotransfected with tau^WT^ and GCaMP6 calcium indicator. Images were taken of GCaMP6 at 2 Hz, showing basal calcium activity within the neuron. TTX was added to silence sodium voltage-gated ion channel mediated action potentials. This video consists of 502 frames and has an image size of 37 x 37 μm. See also Fig. 6b.

**Movie S5. Calcium activity in a tau^E14^ expressing neuron.** This video shows a neuron cotransfected with tau^E14^ and GCaMP6 calcium indicator. Images were taken of GCaMP6 at 2 Hz, showing basal calcium activity within the neuron. TTX was added to silence sodium voltage-gated ion channel mediated action potentials. This video consists of 420 frames and has an image size of 37 x 37 μm. See also Fig. 6c.

